# A clinical mutation in *uvrA*, a DNA repair gene, confers survival advantage to *Mycobacterium tuberculosis* in the host

**DOI:** 10.1101/2024.10.07.616951

**Authors:** Saba Naz, Dipanwita Datta, Sidra Khan, Yogendra Singh, Vinay Kumar Nandicoori, Dhiraj Kumar

## Abstract

DNA repair pathways play an essential role in maintaining the genomic integrity of bacteria, and a perturbation in their biological activity helps bacteria survive under duress. In drug-resistant clinical strains, we identified a Q135K mutation in the *uvrA* gene, a DNA repair pathway gene. To delineate the role of *uvrA* and the Q135K mutation, we generated the gene replacement mutant of UvrA (*RvΔuvrA*) in *Mycobacterium tuberculosis H37Rv* (*Mtb-Rv*). While the lack of UvrA function in *RvΔuvrA* could be restored upon complementation with *uvrA*, the *uvrA-Q135K* mutant identified in clinical drug-resistant strains failed to do so. This was reflected in higher mutation rates in *RvΔuvrA* and *RvΔuvrA::uvrA*_*Q135A*,_ compared with wild-type *Rv* or *RvΔuvrA::uvrA* complemented strains in the presence and absence of oxidative stress. Killing kinetics experiments with anti-TB drugs showed increased survival of *RvΔuvrA* and *RvΔuvrA::uvrA*_*Q135K*,_ strains compared with *Rv* or *RvΔuvrA::uvrA*. Importantly, *RvΔuvrA* and *RvΔuvrA::uvrA*_*Q135K*_ showed enhanced survival in peritoneal macrophages and murine infection model of infection. Together, data suggests that acquiring Q135K mutation benefits the pathogen, which helps enhance the host’s survival adaptability.

**Author Summary:** DNA repair mechanisms in an organism are necessary for correcting the errors generated during replication or when it is damaged/modified due to insults. As a GC organism, Mtb is highly prone to host-mediated attacks on its genome, which, if uncorrected, can impact its genome integrity. The drug-resistant clinical strains of *Mtb* harbor Q135K mutation in *uvrA*, the first enzyme in the nucleotide excision repair pathway. With the help of genetic, molecular, and murine challenge experiments, we show that the UvrA-Q135K mutation abrogates the enzyme’s activity, compromising the *Mtb* strain harboring the mutation in the oxidative and nitrosative stress. On the contrary, the mutation in UvrA imparts survival advantage in activated macrophages and murine infection models. Results presented argue that identified mutation helps in better adaptability in the host, which may include faster acquisition of drug resistance.

## Introduction

*Mycobacterium tuberculosis* (*Mtb*), causes Tuberculosis (TB) and is one of the most ancient pathogens known to humankind. *Mtb*’s ability to stay resilient in a hostile environment is dedicated to the presence of various pathways that help to thwart the killing effects of the host immune system (1). The *Mtb* genome is vulnerable to the impact of the reactive oxygen species and reactive nitrogen intermediates that modify the bases. *Mtb* encodes various DNA repair pathways that maintain its genomic integrity (2). One such pathway is the **<u>n</u>**ucleotide **<u>e</u>**xcision **<u>r</u>**epair pathway (NER), which repairs different lesions (3). NER was first known to repair thymine dimers, and subsequent studies identified that the pathway is activated upon the formation of DNA cross-links, strand breaks, abasic sites, and others. NER is conserved across different bacterial species and is initiated by the action of UvrABC exinuclease. A complex (UvrA)_2_UvrB recognizes the damaged nucleotide in an ATP-dependent manner, followed by the recruitment of a nuclease UvrC. The DNA damage is repaired by the action of enzymes UvrD, DNA polymerase, and DNA ligase (4). Previously, it is reported that the deletion of *uvrA* results in sensitivity towards UV radiation, DNA damaging agents, and redox stress (5). The previous studies indicate the important role of UvrA in *Mtb* biology because this pathogen lacks a mismatch repair pathway (4, 6).

The acquisition of drug resistance in *Mtb* has created an alarming situation. Besides, the emergence of multi and extensively drug-resistant TB has aggravated treatment outcomes (7). Genome sequencing of clinical strains identified direct targets of antibiotics that cause drug resistance (8). However, drug resistance cannot be determined only by the known direct targets of antibiotics, implicating a role of unknown targets responsible for causing drug resistance (9). It is essential to identify novel targets for improved diagnosis in clinical settings. Previous studies have determined drug resistance mechanisms and targets using genome-wide association analysis. *Mtb* has seven major lineages that are spread across the world. Lineage 2 strains tend to acquire drug resistance faster than lineage 4 strains. Moreover, mutations in the DNA repair genes were also found in lineage 2 of *Mtb* (10). Interestingly, we identified mutations in DNA repair genes that belong to the base excision repair pathway, nucleotide excision repair pathway, and homologous recombination using genome-wide association in the MDR/XDR strains (11, 12). We identified a non-synonymous mutation (Q135K) in *uvrA* that was exclusively present in MDR/XDR-TB strain. The findings were intriguing because a previous study showed the vital role of uvrA in combating redox stress. Based on previous findings and analysis, we sought to ask: a) How will Q135K mutation in UvrA impact its function under different stress conditions? b) what is the mutation rate of *RvΔuvrA* and *RvΔuvrA::uvrA*_*Q135A*_ in the presence of oxidative stress drugs? c) Does Q135K have a role to play in the survival and adaptability of *ex vivo* and *in vivo*?

To answer the above questions, we generated and characterized the deletion mutant of *uvrA* in *Mtb*. We performed *in vitro, ex vivo*, and *in vivo* experiments that revealed that the Q135K mutation identified in the *uvrA* abrogates its biological function and helps bacteria resist host stress conditions.

## Result

### Generation and characterization of gene replacement mutant of uvrA in Mtb

To examine the role of UvrA and the identified GWAS mutation in conferring drug resistance, we generated the gene replacement mutant of *uvrA* in *Mtb* using homologous recombination in the laboratory strain *H37Rv* (*Rv*). It is a 2919 bp gene that encodes for ∼106 kDa protein and is non-essential for *Mtb*’s growth *in vitro* and *in vivo*. UvrA has an ATP binding domain and a UvrB interacting domain (1a). The native locus of *uvrA* was disrupted with the hygromycin resistance cassette (Fig 1b & c). Gene replacement mutant of *uvrA* was confirmed with the help of multiple PCRs (Fig 1c). Next, we generated the complementation constructs by cloning wild-type *uvrA* and *uvrA*-Q135K in the integrative vector harboring the FLAG tag at the N-terminus (Fig 1d). The complementation constructs were electroporated in the *uvrA* gene replacement mutant (*RvΔuvrA)*. The expression of *uvrA* and *uvrA*-Q135K was confirmed through western blot analysis. (Fig 1e) We next sought to identify the growth kinetics of the *Rv, RvΔuvrA, RvΔuvrA*::*uvrA*, and *RvΔuvrA*::*uvrA*_*Q135K*_ in 7H9-ADC complete medium. We did not identify any discernible difference in the growth of *Rv, RvΔuvrA, RvΔuvrA*::*uvrA*, and *RvΔuvrA*::*uvrA*_*Q135K*_ (Fig 1f).

**Figure 1:**
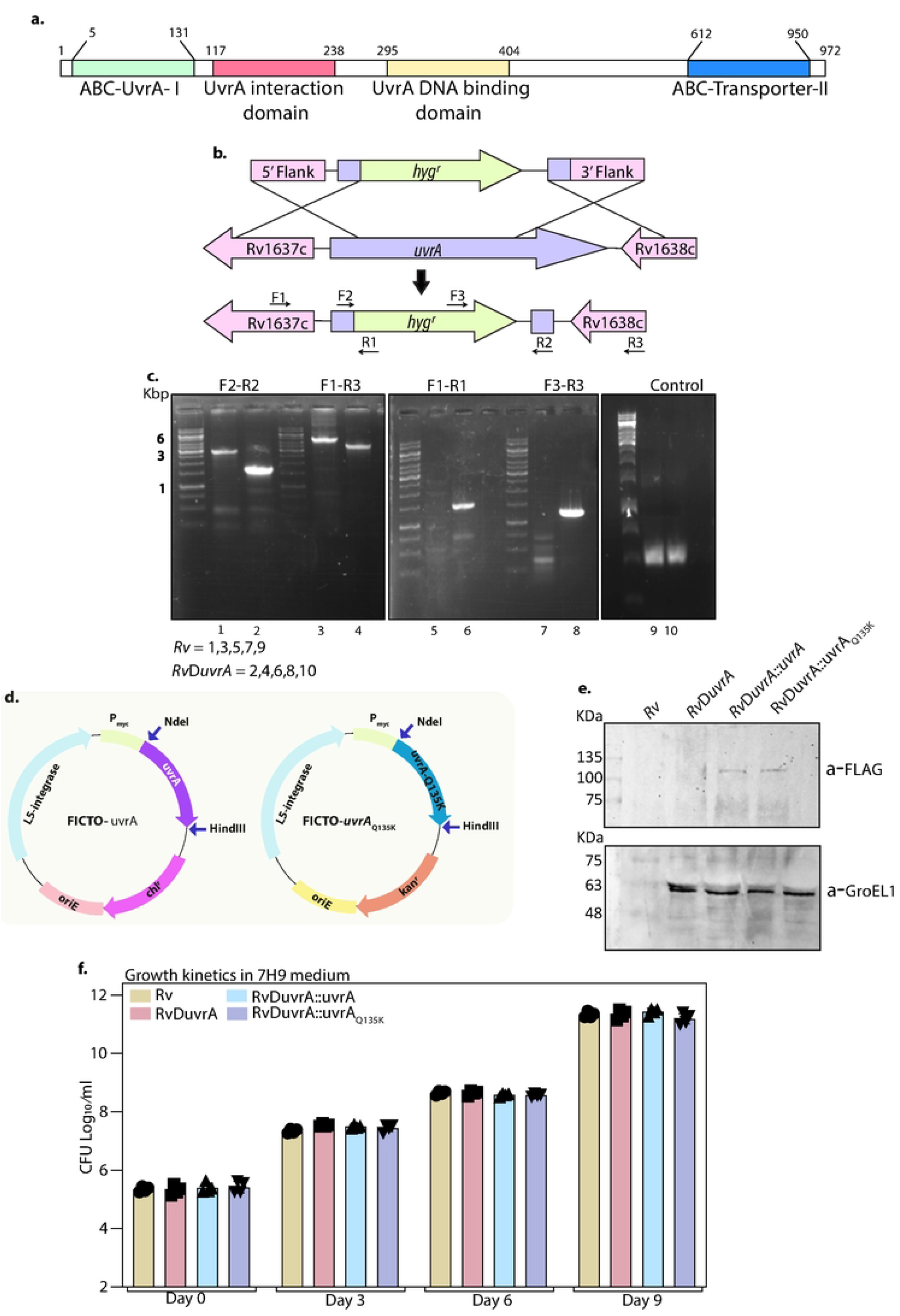
Generation and characterization of gene replacement mutant of uvrA in Mtb. **a)** Schematic depicting the generation of gene replacement mutant of *uvrA*. The *hygromycin*^*r*^ cassette disrupted the native *uvrA* allele. **b)** PCR using F2-R2 (gene-specific primers) amplified 3.1 kb in *Rv* and ∼1.6 kb in *RvΔuvrA*. PCR using F1-R3 amplified a band of ∼6 kb *Rv* and ∼4.6 kb in *RvΔuvrA*. PCR using F1-R1 (*hygromycin*^*r*^ cassette reverse primer and forward primer of the 5’ flank) amplified a band only in *RvΔuvrA*. Similarly, PCR using F3-R3 (*hygromycin*^*r*^ cassette forward primer and reverse primer of the 3’ flank) amplified a band only in *RvΔuvrA. TatA* was performed to ensure an equal amount of genomic DNA in *Rv* and *RvΔuvrA*. **c)** Schematic representing the generation of complementation constructs using the integrative vector. *uvrA-* WT and *uvrA*_Q135K_ were cloned at NdeI and HindIII sites. **d)** Western blot showing the expression of the complementation constructs in the background of *RvΔuvrA* (upper panel) and GroEL-1 as a control (lower panel). **e)** Growth kinetics of *Rv, RvΔuvrA, RvΔuvrA*::*uvrA*, and *RvΔuvrA*::*uvrA* _Q135K_ at days 0, 3,6, and 9.

### RvΔuvrA and RvΔuvrA::uvrA_Q135K_ exhibit compromised growth under stress condition

To investigate the role of UvrA and UvrA-Q235K in modulating the survival of pathogen under stress, we exposed the *Rv, RvΔuvrA, RvΔuvrA*::*uvrA*, and *RvΔuvrA*::*uvrA*_*Q135K*_ to *in vitro* stress conditions such as hypoxia, nitrosative, and oxidative stress. We did not observe any differences in the survival of *Rv, RvΔuvrA, RvΔuvrA*::*uvrA*, and *RvΔuvrA*::*uvrA*_*Q135K*_ under hypoxic conditions (Fig 2a). Compared with *Rv*, we observed a ∼10-fold (1 log_10_ fold) decrease in the survival of *RvΔuvrA*, at 48 h post-addition of sodium nitrite (Fig 2b). While the complementation with *uvrA* (*RvΔuvrA*::*uvrA*) restored the perturbed survival, complementation with *uvrA-Q135K* (*RvΔuvrA*::*uvrA*_*Q135K*_) failed do so, suggesting that Q135K mutation impacts the biological activity of UvrA (Fig 2b). The results were quite similar when the strains *Rv, RvΔuvrA, RvΔuvrA*::*uvrA*, and *RvΔuvrA*::*uvrA*_*Q135K*_ strains were exposed to oxidative stress, wherein we observed ∼9 and 5 fold compromised growth in *RvΔuvrA*, and *RvΔuvrA*::*uvrA*_*Q135K*_ compared with *Rv*, and *RvΔuvrA*::*uvrA* strains. The results were akin to a previous report, in which the absence of *uvrA* showed compromised survival under oxidative stress (5). Together, data suggests that the absence of *uvrA* impacts survival under nitrosative and oxidative stress conditions, and uvrA-Q135A mutation perturbs the enzyme’s function (Fig 2a).

**Figure 2:**
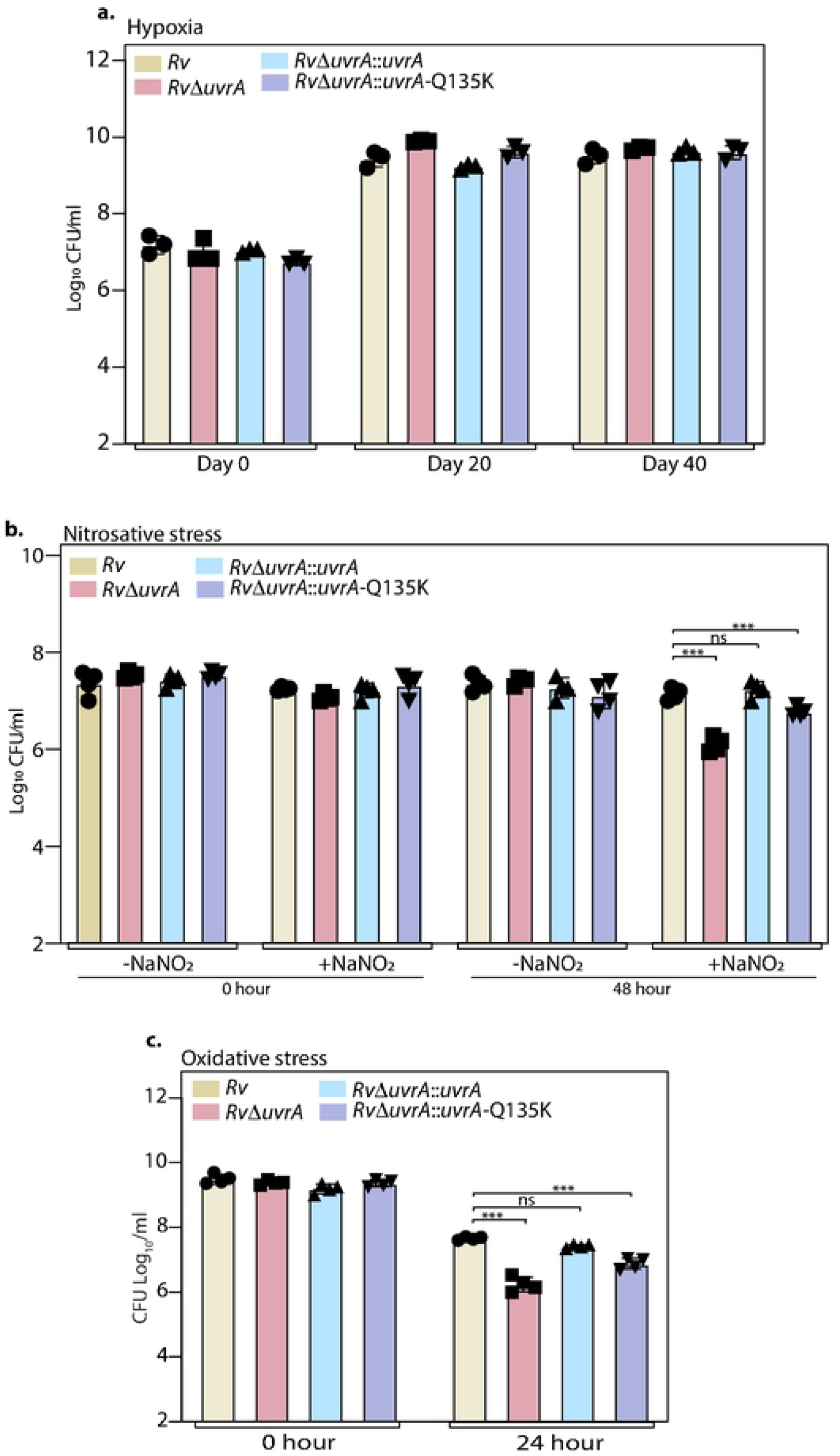
Survival of Rv, RvΔuvrA RvΔuvrA::uvrA, and RvΔuvrA:uvrA_Q135K_ in the presence of in vitro stresses. **a)** Graphs representing the survival of *Rv, RvΔuvrA RvΔuvrA*::*uvrA*, and *RvΔuvrA*:*uvrA*_Q135K_ in the hypoxic conditions at day 0, 20 and 40. **b)** Graph representing the effect of nitrosative stress on *Rv, RvΔuvrA RvΔuvrA*::*uvrA*, and *RvΔuvrA*:*uvrA*_Q135K_ at 0 and 48-hour post addition of sodium nitrite. **c)** Graph representing the survival of oxidative stress at 0 and 24 h post addition of 50 mM CHP. All graphs were generated using the GraphPad prism version 9 and the statistical analysis (two-way ANOVA) was performed using GraphPad prism software. *** p<0.0001, **p<0.001 and *p<0.01.

### RvΔuvrA and RvΔuvrA::uvrA_Q135K_ strains show higher mutation rates in the absence and presence of oxidative stress

UvrA initiates the nucleotide excision repair pathway with the help of UvrB. Upon binding the DNA, the damaged nucleotide is excised that subsequently recruits UvrC, a nuclease, that excises 12 or 13 nucleotides. Next, UvrD, a DNA helicase, binds and disassembles the DNA, leaving a gap filled by the DNA polymerase I and sealed by the DNA ligase (Fig 3a) (3). Since UvrA belongs to the DNA repair pathway, we sought to investigate the mutation rate of *Rv, RvΔuvrA, RvΔuvrA*::*uvrA*, and *RvΔuvrA*::*uvrA*_*Q135K*_ strains with the help of David fluctuation assay (Fig 3b)(13). The spontaneous mutation rate of *RvΔuvrA*, and *RvΔuvrA*::*uvrA*_*Q135K*_ strains spontaneous was higher than *Rv*, and *RvΔuvrA*::*uvrA* when the cultures were exposed to ciprofloxacin, isoniazid and rifampicin. *RvΔuvrA*, and *RvΔuvrA*::*uvrA*_*Q135K*_ showed ∼2.65 and 2.38-fold increase in the spontaneous mutation rate in the presence of ciprofloxacin (Fig 3c). Similarly, the mutation rate of *RvΔuvrA*, and *RvΔuvrA*::*uvrA*_*Q135K*_ was 2.04 and 2.37 in the presence of isoniazid and 4.27 and 3.68, in the presence of rifampicin (Fig 3d-f).

**Figure 3:**
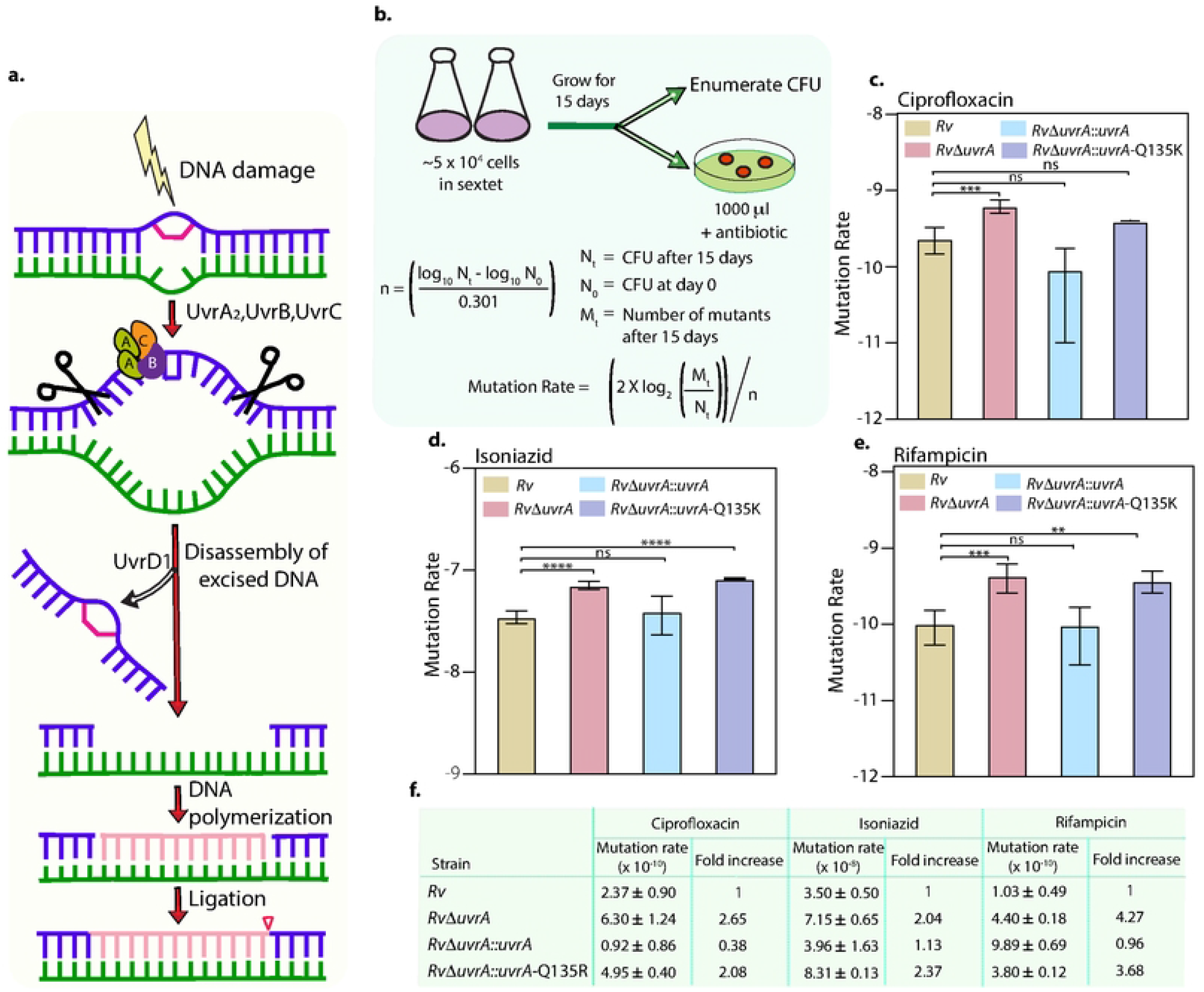
Mutation rate analysis. **a)** Schematic representing the initiation of the NER pathway upon DNA damage. **b)** Mutation rate analysis using the David fluctuation assay. **c-e)** Mutation rate analysis of *Rv, RvΔuvrA RvΔuvrA*::*uvrA*, and *RvΔuvrA*:*uvrA*_Q135K_ in the presence of ciprofloxacin, isoniazid and rifampicin. **f)** Fold change of *Rv, RvΔuvrA RvΔuvrA*::*uvrA*, and *RvΔuvrA*:*uvrA*_Q135K_ in the presence of ciprofloxacin, isoniazid and rifampicin. All graphs were generated using GraphPad Prism version 9 and the statistical analysis (two-way ANOVA) was performed using GraphPad Prism software. *** p<0.0001, **p<0.001 and *p<0.01.

One of the contributors to the DNA damage in bacteria or the host is the oxidative stress. The inability to neutralize the reactive intermediates and the reactive oxygen species (ROS) causes oxidative stress (14). We observed a log_10_-fold decrease in the survival of the *RvΔuvrA*, and *RvΔuvrA*::*uvrA*_*Q135K*_ in the presence of oxidative stress (Fig 2c), therefore, we sought to investigate the spontaneous mutation rate in the presence of oxidative stress by with the help of David fluctuation assay, in which the strains were subjected for 24 h prior to plating to CHP treatment. *Rv, RvΔuvrA, RvΔuvrA*::*uvrA*, and *RvΔuvrA*::*uvrA*_*Q135K*_ strains were grown in the 7H9-ADC medium for 14 days, and subsequently CHP was added for 24 h and mutation rate was determined (Fig 4a). The mutation rate observed in case of *RvΔuvrA* and *RvΔuvrA*::*uvrA*_*Q135K*_ was 9.7 and 6.4 fold compared with the *Rv* and *RvΔuvrA::uvrA*_*Q135K*_ in the presence of ciprofloxacin (Fig 4b&e). Similarly, we observed 63.04 and 13.69 fold increase in the mutation rate of the *RvΔuvrA* and *RvΔuvrA*::*uvrA*_*Q135K*_ compared with the *Rv* and *RvΔuvrA::uvrA* in the presence of isoniazid (Fig 4c & e). The mutation rate observed in case of *RvΔuvrA* and *RvΔuvrA*::*uvrA*_*Q135K*_ was 59.7 and 22.33 fold higher than the *Rv* and *Rv*Δ*uvrA*::*uvrA* in the presence of rifampicin (Fig 4d & e). The results suggests that the Q135K mutation in UvrA abrogates its function (Fig 3 &4). Higher mutation rates observed in the absence of UvrA or upon complementation with UvrA-Q135K mutant is suggestive of the ability of these strains to acquire advantageous mutations when subjected to stress conditions.

**Figure 4:**
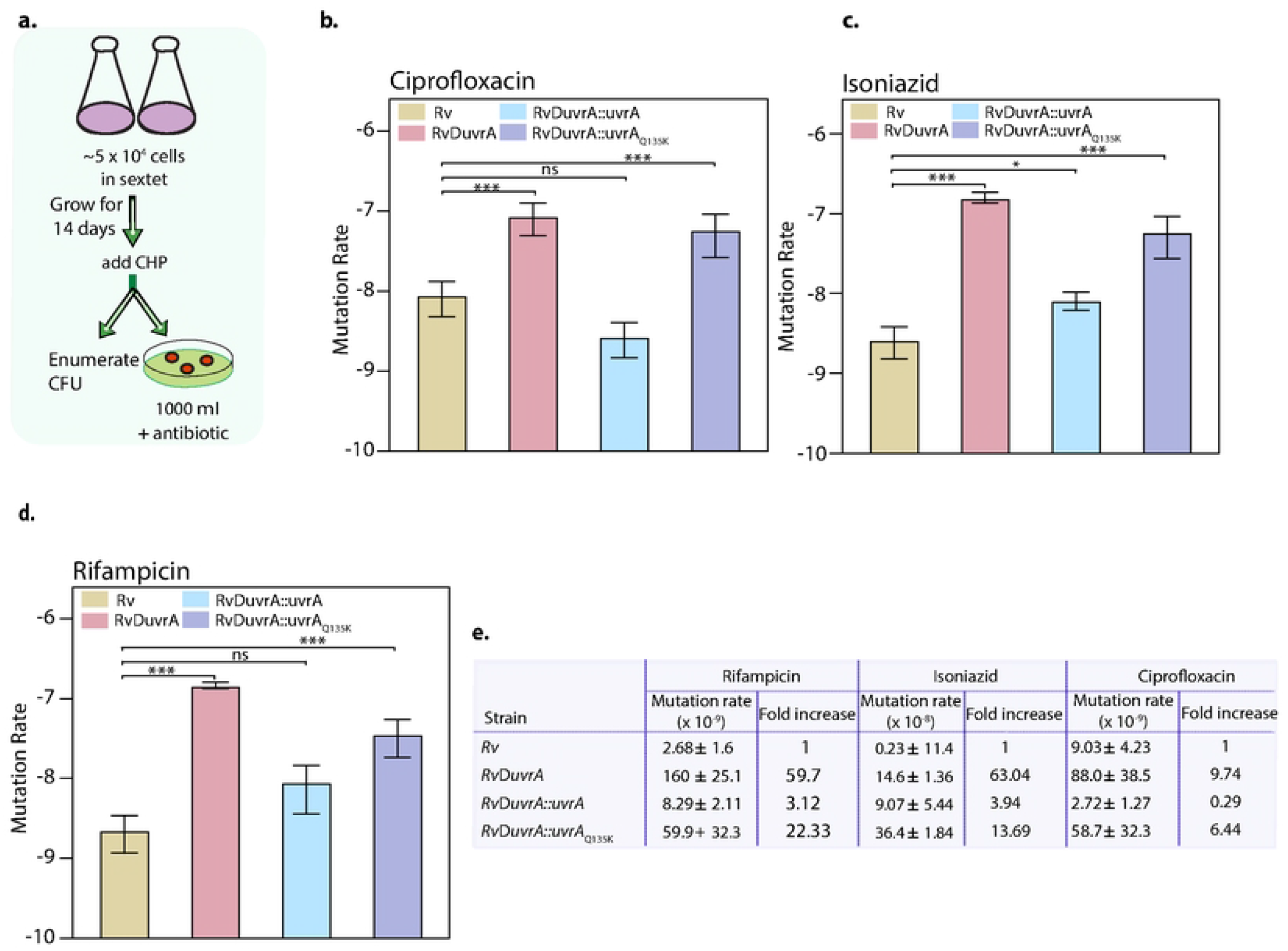
Mutation rate analysis in the presence of oxidative stress. **a)** Schematic representing the mutation rate analysis in the presence of CHP. **b-d)** Mutation rate analysis of *Rv, RvΔuvrA RvΔuvrA*::*uvrA*, and *RvΔuvrA*:*uvrA*_Q135K_ in the presence of ciprofloxacin, isoniazid and rifampicin after CHP treatment. **e)** Fold change of *Rv, RvΔuvrA RvΔuvrA*::*uvrA*, and *RvΔuvrA*:*uvrA*_Q135K_ in the presence of ciprofloxacin, isoniazid and rifampicin. All graphs were generated using the GraphPad prism version 9 and the statistical analysis (two-way ANOVA) was performed using GraphPad prism software. *** p<0.0001, **p<0.001 and *p<0.01.

### *RvΔuvrA* and *RvΔuvrA*::*uvrA*_*Q135K*_ strains show increased survival in the presence of antibiotics

We next investigated the killing kinetics of *Rv, RvΔuvrA, RvΔuvrA*::*uvrA*, and *RvΔuvrA*::*uvrA*_*Q135K*_ in the absence and presence of different anti-TB drugs. In line with the results in Fig 1, in the absence of antibiotics, we did not observe a significant difference in the survival between the strains (Fig 5a). As expected, when the growing strains were exposed to anti-TB drugs, they showed compromised survival from day 3, and the difference was more than 2 log_10_ fold after 9 days of treatment (Fig 5b-d). While death was observed upon anti-TB drug treatment for all strains, in comparison with *Rv, RvΔuvrA, RvΔuvrA*::*uvrA*, and *RvΔuvrA*::*uvrA*_*Q135K*_ strains showed better survival (Fig 5b-d). The survival of *RvΔuvrA*, and *RvΔuvrA*::*uvrA*_*Q135K*_ strains were ∼0.5 log_10_ fold higher, suggesting that the absence of UvrA or having Q135K mutation is likely to help in combating anti-TB drugs. This could be due to the acquisition of mutations due to compromised repair activity, which may be contributing towards improved ability to combat anti-TB drug treatment.

**Figure 5:**
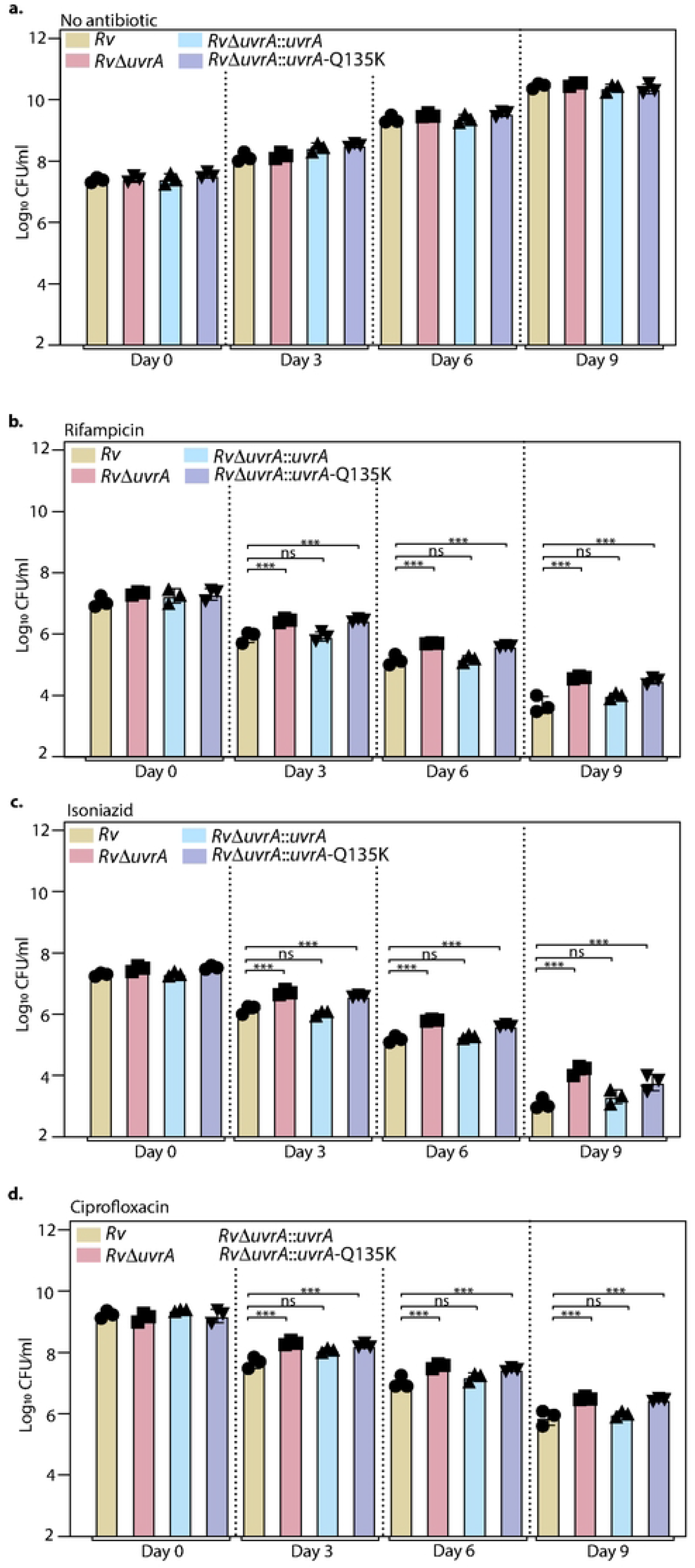
Killing kinetics in the presence of anti-TB drugs. **a)** Survival of *Rv, RvΔuvrA RvΔuvrA*::*uvrA*, and *RvΔuvrA*:*uvrA*_Q135K_ without antibiotics. **b-d)** Survival of *Rv, RvΔuvrA RvΔuvrA*::*uvrA*, and *RvΔuvrA*:*uvrA*_Q135K_ in the presence of rifampicin, isoniazid and ciprofloxacin. Different anti-TB drugs were added on day 0, 3, 6 and 9. All graphs were generated using GraphPad Prism version 9 and the statistical analysis (two-way ANOVA) was performed using GraphPad Prism software. *** p<0.0001, **p<0.001 and *p<0.01.

### *RvΔuvrA* and *RvΔuvrA*::*uvrA* Q135K resist the killing effects of antibiotics *ex vivo*

Type II interferon plays many roles, including activating macrophages, promoting bactericidal activity, and stimulating phagocytosis (15). To determine the impact of interferon-gamma (IFN-γ) on the survival of the *Rv, RvΔuvrA, RvΔuvrA*::*uvrA*, and *RvΔuvrA*::*uvrA*_Q135K_, we performed *ex vivo* infection experiments. Peritoneal macrophages were stimulated with IFN-γ, and different anti-TB drugs were added at 24 hours post-infection (h.p.i), and CFUs were determined (Fig 6a). At 4 and 24 h.p.i, we did not observe any differences in survival (Fig 6b). At 48 h.p.i of IFN-γ addition, we observed a 10-fold increase in the survival of *RvΔuvrA* compared with the IFN-γ untreated set (Fig 6b). This result is congruent with the recently published study wherein the *uvrA* mutant exhibits ∼15 to 20-fold increase in survival in the bone marrow-derived macrophages (BMDM) when stimulated with IFN-γ (16). Moreover, a ∼5-fold increase in the survival of *RvΔuvrA*::*uvrA*_Q135K_ was observed, suggesting that the Q135K mutation in UvrA has affected its biological function (Fig 6). Next, we determined the survival of *Rv, RvΔuvrA, RvΔuvrA*::*uvrA*, and *RvΔuvrA*::*uvrA*_Q135K_ in the presence of anti-TB drugs with and without IFN-γ. In the presence of isoniazid, ciprofloxacin, and rifampicin, an increase of ∼5, 3, and 5-fold in the survival of *RvΔuvrA* was observed (Fig 6c). In the presence of IFN-γ and isoniazid, ciprofloxacin, and rifampicin, ∼20, 15, and 20-fold increase in the CFUs of *RvΔuvrA* (Fig 6c). The data suggest that IFN-γ and anti-TB drugs have a synergistic effect on the survival of *RvΔuvrA*. Furthermore, *RvΔuvrA*::*uvrA* restores the phenotype, whereas *RvΔuvrA*::*uvrA*_Q135K_ showed a similar trend as *RvΔuvrA* (Fig 6c). The findings indicate host stress and anti-TB drugs are the essential drivers in promoting the survival of *RvΔuvrA* and *RvΔuvrA*::*uvrA*_Q135K_ (Fig 6).

**Figure 6:**
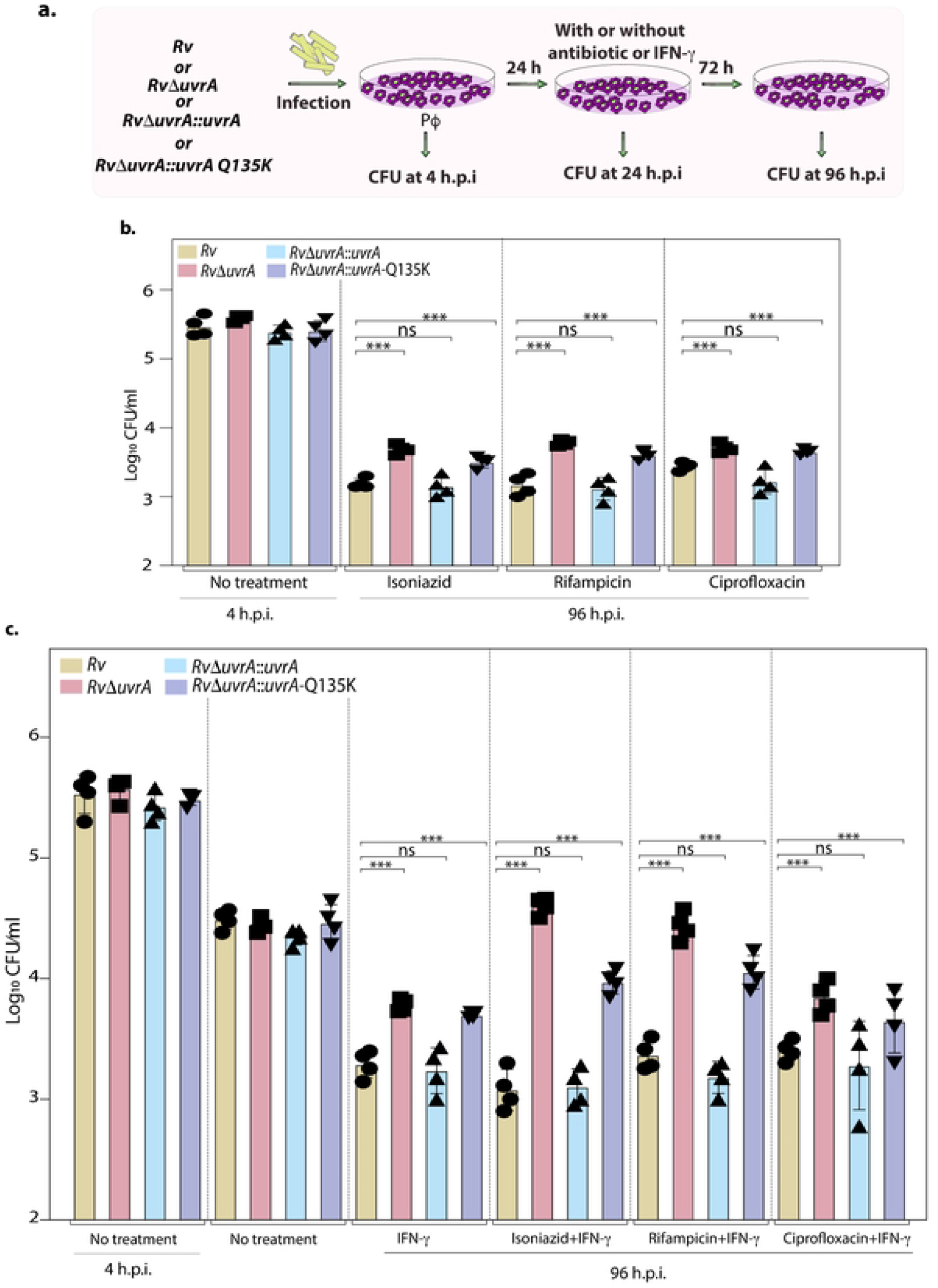
Ex vivo survival of Rv, RvΔuvrA, RvΔuvrA::uvrA and RvΔuvrA::uvrA-Q135K. **a)** Schematic representing the *ex vivo* infection experiment in the peritoneal macrophages. Cells were infected with *Rv, RvΔuvrA RvΔuvrA*::*uvrA*, and *RvΔuvrA*:*uvrA*_Q135K_. CFUs were enumerated at 4 h.p.i. to determine the uptake. Infected cells were either not treated or treated with the IFN-γ, isoniazid, rifampicin, and ciprofloxacin independently or in combination. CFUs were enumerated 96 h.p.i. **b-c)** Graphs representing the CFUs at different time points. All graphs were generated using the GraphPad prism version 9 and the statistical analysis (two-way ANOVA) was performed using GraphPad prism software. *** p<0.0001, **p<0.001 and *p<0.01.

### *RvΔuvrA* and *RvΔuvrA*::*uvrA* Q135K exhibit better survival in the host

As observed in the above *ex vivo* survival, host stress plays a crucial role in the growth of the pathogen during infection (17). On one hand, in the absence of important genes, bacteria show compromised growth in the host stress. On the other hand, a consistent increase in the survival of the DNA repair deficient strains was observed in the host (18). We performed murine infection experiments to determine the growth phenotype of the *RvΔuvrA* and *RvΔuvrA::uvrA*_*Q135K*_ in the host (Fig 7a). We challenged the mice through an aerosol route and determined the CFU 1-day and 8 weeks post-infection. No significant difference in the day-1 CFU was observed in *Rv, RvΔuvrA, RvΔuvrA::uvrA*, and *RvΔuvrA*::*uvrA*_Q135K,_ suggesting the equivalent implantation of different strains in the mice lungs (Fig 7b). At four weeks, we observed a ∼5-fold increase in the survival of *RvΔuvrA*, and *RvΔuvrA*::*uvrA*_Q135K_ in the lungs and a ∼2-3-fold increase in the survival in the spleen compared to *Rv* and *RvΔuvrA::uvrA*. The difference was more apparent in the 8-week post-infection. A log fold increase was observed in the survival of *RvΔuvrA*, and *RvΔuvrA*::*uvrA*_Q135K_ compared to *Rv* and *RvΔuvrA::uvrA*. Collectively, the data suggest that the deletion of *uvrA* from *Mtb* helps in better survival in the mice, and Q135K abrogates the function of UvrA (Fig 7).

**Figure 7:**
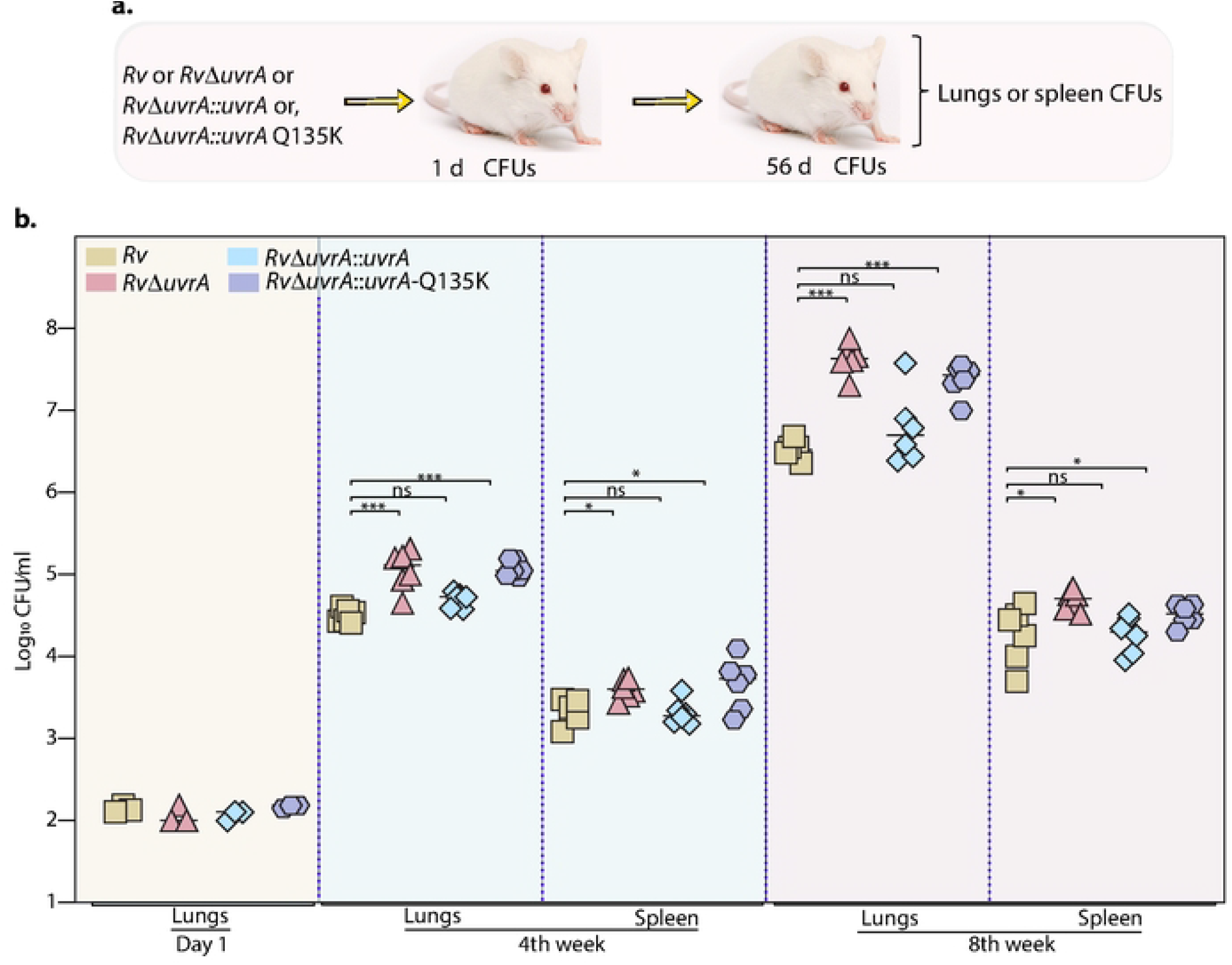
In vivo survival of Rv, RvΔuvrA, RvΔuvrA::uvrA and RvΔuvrA::uvrA-Q135K. **a)** Schematic representing the *in vivo* infection experiment in the mice. Infection was performed using *Rv, RvΔuvrA, RvΔuvrA*::*uvrA*, and *RvΔuvrA*::*uvrA* _Q135K_ through aerosol. Lungs were isolated at day 1, 4^th^ week, and 8^th^ week post-infection for CFU analysis. **b)** Graph showing the CFU analysis of the lungs and spleen at the 4^th^ and 8^th^ week post-infection. All graphs were generated using the GraphPad Prism version 9 and the statistical analysis (two-way ANOVA) was performed using GraphPad Prism software. *** p<0.0001, **p<0.001 and *p<0.01.

## Discussion

Drug resistance in *Mtb* has significantly impacted the treatment outcome of TB. The emergence of **<u>m</u>**ulti**<u>-d</u>**rug **<u>r</u>**esistant (MDR-TB) and e**<u>x</u>**tensively **<u>d</u>**rug**<u>-r</u>**esistant TB (XDR-TB) is now a major global concern (19). Detection of the drug-resistant -TB is crucial for treating the disease at early onset (20). While efforts are dedicated to detecting resistance in clinical settings using molecular tools, the involvement of gene mutations classically not known to confer drug resistance makes the detection more challenging (9). The resistance against standard-of-care anti-TB drugs is typically driven by the genes involved in central metabolic pathways, cell-wall biosynthesis, DNA metabolism, etc. Surprisingly, clinically, all drug resistance is not attributable to these select genes (21). Similarly, clinical isolates showing resistance to multiple drugs do not often show mutations in all of the corresponding classical target genes (22). Therefore, identifying mechanisms that could contribute to resistance against various drugs is critical to developing a better predictive tool for drug resistance. Multiple studies identified and validated novel targets contributing to drug resistance in clinical settings. For example, the experimental evolution of drug resistance against bedaquiline and clofazimine showed a strong association with the transcriptional repressor Rv0678 (23).

Similarly, an analysis of 51,229 clinical isolates of *Mtb* identified a transcriptional regulator, *Rv1830*, that impacts the effect of antibiotic post-treatment. Rv1830, with other transcriptional factors, controls cell growth and division. Mutations in *Rv1830* are associated with ineffective treatment and drug resistance (24). In addition to the transcription factors discussed above, mutations in DNA repair pathway genes are associated with drug resistance (25-28). Lineage 2 strains, like Beijing family strains, have high inclination towards acquiring drug resistance (29). Genome sequence analysis of Beijing family strains identified polymorphism in the DNA repair pathway genes *mutT2, mutT4*, and *ogt* (26, 27). Moreover, mutations in the *dnaA*, and *dnaQ* (DNA replication), *alkA* and *nth* (base excision repair pathway) and *recF* (homologous recombination pathway) were also reported in drug-resistant strains (28). However, without any mechanistic study, it was proposed that the observed mutations in the DNA repair genes could be due to random mutations or hitchhiking (26, 28). Therefore, bacterial proteins involved in genome-scale functions, like transcriptional regulation, DNA repair, etc., acquire significance in understanding the mechanisms of drug resistance and developing potential markers for early diagnosis of drug-resistant TB.

Previously, we showed that the deletion mutant of *ung, udgB*, and *RvΔung-udgB* showed better survival in the peritoneal macrophages and guinea pig lungs (11). Genome sequencing of the *RvΔung-udgB* isolated from guinea pig lungs shows the acquisition of mutations across the genome, suggesting that host pressure is a critical determinant in shifting the equilibrium towards selecting bacterial strains that can adapt to the host environment (11). Reactive nitrogen intermediates, reactive oxygen species, and immune system modulators like cytokines trigger the induction of stress response pathways that aid bacteria in enduring the killing effects of antibiotics. In the case of the DNA repair deficient strains, the stress response and evolution are faster due to the hypermutable phenotype (17, 21). In a subsequent genome-wide association analysis to identify the genetic triggers that aid in acquiring drug resistance in clinical settings, we identified mutations in the DNA repair pathway genes. *MutY* belongs to the base excision repair pathway and was mutated in the drug-resistant strains. Experiments showed that the *mutY* deletion strain harboring the alternative allele of *mutY* failed to rescue the phenotype of *mutY* (12).

In line with the previous findings, we determined the effect of Q135K mutation in *uvrA*. Using *in vitro, ex vivo*, and *in vivo* infection experiments, we identified that gene replacement mutant of *uvrA* and *RvΔuvrA*::*uvrA*_Q135K_ exhibits better adaptability in the host. In conclusion, the study reports the role of clinically relevant mutation Q135K in providing survival advantage to bacteria. Moreover, Q135K mutation in *uvrA* may be employed as a marker for the detection of drug resistance in clinical settings.

## Contribution

S.N. conceptualized the study. D.K., V.K.N, and Y.S. supervised the study. S.N. and DK acquired the funding. S.N. performed all characterization, *in vitro, ex vivo, and in vivo* infection experiments. S.K. generated gene replacement mutants. S.N, D.D performed *ex vivo* and mice animal infection. S.N. generated figures and written manuscripts. S.N., V.K.N & D.K. review and edited the manuscript.

## Acknowledgment

This work is partly funded by the Ben Barres Spotlight Award funding to S.N. and by the Department of Biotechnology, Government of India (BT/IC-06/003/91-Flagship Program to D.K. We thank the Tuberculosis Aerosol Challenge Facility at ICGEB and its staff for their help in performing animal infection experiments. Vector-pYUB1471 is a kind gift from Prof. William R. Jacobs’s laboratory.

## Competing interest

The authors declare no competing interest.

## Ethics

The animal experiments protocol was approved by the Animal Ethics Committee of the International Centre for Genetic Engineering and Biotechnology, New Delhi, India. The approval (Ethics approval: ICGEB/IAEC/22092022/CI-27) is as per the guidelines issued by the Committee for the Purpose of Control and Supervision of Experiments on Animals (CPCSEA), Government of India.

## Materials and Methods

### Generation of gene replacement mutant and complementation constructs

*UvrA* gene replacement mutant was generated using the recombineering methods. The upstream and downstream regions of *uvrA* were amplified using the primers and the amplified products were digested using PflM1. Separately, the pYUB1471 vector was digested with PflM1 to obtain the *oriE*+ *λ cos* sites DNA fragment (30). The three digested fragments were ligated with the hygromycin-resistant cassette. The allelic exchange substrate thus obtained was digested with the SnaBI to generate a linearized substrate for electroporation in the pNIT-ET overexpressing strain (31). Recombinant colonies were screened using the multiple set of primers for PCR reaction. *uvrA-*WT and *uvrA*_Q135K_ were PCR amplified and digested with NdeI and HindIII. The digested amplicons were cloned in integrative vector FICTO. The complementation constructs were electroporated in the *RvΔuvrA* background to generate *RvΔuvrA*::*uvrA* and *RvΔuvrA*::*uvrA*_Q135K_.

### Western Blot

30 ml cultures of *Rv, RvΔuvrA RvΔuvrA*::*uvrA*, and *RvΔuvrA*:*uvrA*_Q135K_ were inoculated in 7H9-ADC medium at A_600_ ∼0.1 and grown till A_600_ ∼0.8. After pelleting cells in 50 ml falcon tubes at 4000 rpm for 10 min, cells were resuspended in the lysis buffer (1X PBSG) containing protease inhibitors. Resuspended cells were transferred in the zirconium beads containing bead-beating tubes. Bead beating was performed for 6 cycles. The cell lysate was centrifuged twice at 13000 rpm for 45 min at 4^°^C to obtain. The Bradford assay reagent was used for the protein estimation. 50 μg of *Rv, RvΔuvrA RvΔuvrA*::*uvrA*, and *RvΔuvrA*:*uvrA*_Q135K_ were loaded on 10% SDS-PAGE independently and transferred to nitrocellulose membrane. The membrane was blocked using 5% BSA prepared in 1XPBST_20_ for 2 h. a-FLAG (1:5000), and a-GroEL-1 (1:10000) were incubated with membranes overnight at 4^°^C. Membranes were washed using 1XPBST_20_ (thrice) and incubated with anti-rabbit secondary antibody DARPO (1:10000) or anti-mouse secondary antibody for 2 h at room temperature (25^°^C). After washing membranes thrice with 1XPBST_20,_ blots were developed using a chemiluminescence reagent.

### Growth kinetics and *in vitro* stress

For *in vitro* growth analysis, *Rv, RvΔuvrA RvΔuvrA*::*uvrA*, and *RvΔuvrA*:*uvrA*_Q135K_ were grown in 7H9-ADC medium up to A_600_∼0.6 and inoculated into a fresh medium at the final of A_600_∼0.1 in biological triplicates. CFUs were enumerated at days 0, 3, 6, and 9 on 7H11-OADC plates. For oxidative stress *Rv, RvΔuvrA RvΔuvrA*::*uvrA*, and *RvΔuvrA*:*uvrA*_Q135K_ were inoculated at A_600_∼0.1 in 7H9-ADS medium and treated with 50 mM CHP. CFUs were enumerated at 0 and 24 h post-CHP addition on 7H11-OADC-containing plates. For nitrosative stress, *Rv, RvΔuvrA RvΔuvrA*::*uvrA*, and *RvΔuvrA*:*uvrA*_Q135K_ were inoculated at A_600_∼0.2 in 7H9-ADC medium, pH=5.5 and treated with a final concentration of 3 mM sodium nitrite. CFUs were enumerated at 0 and 48 h post-treatment. For hypoxia, *Rv, RvΔuvrA RvΔuvrA*::*uvrA*, and *RvΔuvrA*:*uvrA*_Q135K_ were inoculated at A_600_∼0.1 in 7H9-ADC medium. Cultures were added in the cryovials (95% head space ratio) and sealed with parafilm. CFUs were enumerated at days 0, 20, and 40 post-hypoxia on 7H11-OADC-containing plates. Experiments were performed using two independent biological experiments and each biological experiment was performed in triplicates. Statistical analysis (one-way ANOVA) was performed using n=4 for each biological experiment. GraphPad Prism software is used for statistical analysis. *** p<0.0001, **p<0.001 and *p<0.01.

### Mutation rate analysis

Mutation rate analysis was performed as described previously (32). Briefly, 50000 cells of *Rv, RvΔuvrA RvΔuvrA*::*uvrA*, and *RvΔuvrA*:*uvrA*_Q135K_ were inoculated in 7H9-ADC medium containing 15% sterile culture supernatant of *Rv. Rv, RvΔuvrA RvΔuvrA*::*uvrA*, and *RvΔuvrA*:*uvrA*_Q135K_ were grown in sextet for 15 days. On the 15^th^ day plating was performed on plain 7H11-OADC plates and isoniazid (5 *μ*g/ml), rifampicin (2 *μ*g/ml), and ciprofloxacin (1.5 *μ*g/ml) containing 7H11-OADC plates. For determining mutation rate in the presence of oxidative stress, the same protocol was followed and on the 14^th^ day, cultures were treated with 50 *μ*M CHP and plating was performed on plain 7H11-OADC plates, and isoniazid (1 *μ*g/ml), rifampicin (2 *μ*g/ml) and ciprofloxacin (3 *μ*g/ml) containing 7H11-OADC plates. Each experiment was performed in sextet. Statistical analysis (one-way ANOVA) was performed using n=6. GraphPad Prism software was used for statistical analysis. *** p<0.0001, **p<0.001 and *p<0.01.

### Killing kinetics

*Rv, RvΔuvrA RvΔuvrA*::*uvrA, and RvΔuvrA*:*uvrA*_Q135K_ were inoculated in 7H9-ADC medium at A_600_∼0.1. Cultures were either treated with isoniazid (5 *μ*g/ml), rifampicin (2 *μ*g/ml), and ciprofloxacin (1.5 *μ*g/ml) or not treated with drugs. CFUs of different sets were enumerated on days 0, 3, 6, and 9. Each experiment was performed in quadruplet. Statistical analysis (one-way ANOVA) was performed using n=4. GraphPad Prism software was used for statistical analysis. *** p<0.0001, **p<0.001 and *p<0.01.

### *Ex vivo* infection experiments

Thioglycollate was injected in Balb/c mice, and 120 h post-injection, peritoneal macrophages were isolated. 0.1 million cells were seeded in a 48-well plate. Cells were infected with *Rv, RvΔuvrA RvΔuvrA*::*uvrA*, and *RvΔuvrA*:*uvrA*_Q135K_ independently at an MOI of 1:5. After 24 h p.i cells were treated with IFN-γ (100 units per well), rifampicin (1*μ*g/ml), isoniazid (1*μ*g/ml) and ciprofloxacin (2.5 *μ*g/ml), IFN-γ + isoniazid, IFN-γ + rifampicin, and IFN-γ + ciprofloxacin. CFUs were enumerated at 4 h.p.i and 24 h.p.i. Finally, Cells were lysed at 120 h p.i. using 0.05% SDS, for CFU enumeration on 7H11-OADC containing plates. The experiment was performed in a biological quadruplicate. Statistical analysis (two-way ANOVA) was performed using GraphPad Prism software. *** p<0.0001, **p<0.001 and *p<0.01.

### *In vivo* infection experiment

*Rv, RvΔuvrA RvΔuvrA*::*uvrA, and RvΔuvrA*:*uvrA*_Q135K_ cultures were grown in 7H9-ADC medium up to A_600_ ∼0.8. Cultures were pelleted at 4000 rpm at room temperature (RT). Cells were resuspended in saline and passed through a 26 1/2 gauge needle to obtain a single-cell suspension. 2 × 10^8^ cells were taken in the 15 ml saline for infection. Female Balb/c mice were challenged using a Madison chamber calibrated to deliver ∼ 200 bacilli/lung through the aersolic route. CFUs in the lungs were enumerated at one day post-infection on 7H11-plain plates for *Rv, RvΔuvrA RvΔuvrA*::*uvrA*, and *RvΔuvrA*:*uvrA*_Q135K_ (n=3 for each strain). Survival of *Rv, RvΔuvrA RvΔuvrA*::*uvrA*, and *RvΔuvrA*:*uvrA*_Q135K_ in the lungs and spleen was determined at 4^th^ (n=6 for each strain) and 8^th^-week post-infection (n=6 for each strain). Statistical analysis (two-way ANOVA) was performed using GraphPad Prism software. *** p<0.0001, **p<0.001 and *p<0.01.

## Notes

### Competing Interest Statement

The authors have declared no competing interest.

